# Integration of mechanical testing, *in vivo* optical coherence elastography and personalized finite element modeling to predict geometrical outcomes of corneal cross-linking

**DOI:** 10.1101/2025.09.02.673622

**Authors:** Matteo Frigelli, Robert Lohmüller, Miguel A. Ariza Gracia, Günther Schlunck, Stefan J. Lang, Emilio A. Torres-Netto, Farhad Hafezi, Philippe Büchler, Sabine Kling

## Abstract

**Purpose:** Corneal cross-linking (CXL) induces both mechanical and geometrical changes in the cornea, which are typically overlooked in pre-operative planning. We propose a patient-specific finite element model (FEM) to predict the topographic alterations resulting from CXL.

**Methods:** To calibrate the model, we performed nanoindentation and *ex vivo* OCE inflation tests before and after CXL on five human donor corneas. Nanoindentation results tuned the visco-hyperelastic parameters, while *ex vivo* OCE axial strains were used for validation. Personalized corneal models were generated from the topographies of three keratoconus patients, with regional stiffness in the affected areas adjusted based on axial strain measured by *in vivo* pressure-modulated OCE. Simulated CXL outcomes were then compared to 6-month clinical results.

**Results:** CXL induces a 16-fold increase in the fiber-related mechanical parameters and reduces the viscoelasticity time constant by three. *In vivo* OCE measurements showed an average mechanical weakening of 57% in the KC regions. When compared to the clinical topography at the 6-month follow-up, the CXL-induced curvature changes predicted by the model were −1.5 D vs. −1.76 D, −1.65 D vs. - 1.91 D, and −1.76 D vs. −1.57 D, for the three patients, respectively.

**Conclusion:** By combining FEM with *in vivo* corneal mechanical characterization, patient-specific topographic changes can be predicted, which can be used to improve the planning of CXL treatments.

## Introduction

Keratoconus (KC) is a progressive eye disease in which the cornea, the clear front surface of the eye, thins and bulges outwards in a cone shape. In patients with keratoconus (KC), the cornea exhibits an area where the thickness substantially decreases and the curvature reaches exceptionally high values [1]. This leads to vision impairments and, in severe cases, necessitates corneal transplantation [2]. Localized biomechanical degeneration is a hallmark of this disease [3]. This has been corroborated by x-ray scattering measurements showing a modified collagen orientation and distribution in the KC region [4] with a disarrangement of the orthogonal organization of the fibers [5]. Since the unique distribution of stromal collagen fibers is key for maintaining corneal shape and biomechanics [6], alteration to this organization leads to biomechanical impairment. Accordingly, a reduction in elastic modulus was found in KC corneas compared to healthy controls, measured either *ex vivo* or *in vivo* using uniaxial tensile testing (UTT) [7,8], Brillouin microscopy [9,10], acoustic radiation force elastic microscopy [11], air puff [12,13], nanoindentation (NI) [14], and optical coherence elastography (OCE) [15]. This mechanical weakening and fiber disorganization results in tissue thinning and bulging under the constant intraocular pressure (IOP).

Ultraviolet-A (UVA) corneal cross-linking (CXL) is a photochemical procedure that aims at stiffening the cornea by creating cross-links in the extracellular matrix [16], and is a long-established method of slowing and halting the progression of KC [17]. The original “Dresden protocol” consists of removing the central 8–9 mm of the corneal epithelium, saturating the stroma with riboflavin, and then irradiating the anterior stroma with UVA light (370 nm, 3 mW/cm^2^) for 30 minutes [18]. New CXL protocols are currently under development, aiming at locazing the procedure on the KC, thus increasing the stiffening effect on the weaker region of the cornea. In this context, the ELZA-photo-therapeutic keratectomy (PTK)-assisted customized epi-on (ELZA-PACE) CXL protocol was developed. This novel CXL method relies on increasing riboflavin absorption and delivering higher energy to the KC region to enhance the CXL effect in the targeted area.

CXL has been shown to not only stiffen the cornea, but also to partially reverse the steepening of the cone by flattening the corneal topography [19]. However, the efficacy and the nature of the CXL-induced refractive changes are not completely understood and are therefore not considered in CXL planning. To predict the outcome of CXL in terms of refractive correction, a better understanding of the interplay between the degree of tissue stiffening and the resulting refractive change is required [20]. The mechanical effects of CXL have been extensively described in recent years, with several *ex vivo* studies reporting an increase in the elastic modulus in human corneas after CXL, assessed using mechanical testing techniques such as UTT [21], NI [22], atomic force microscopy [23] and OCE [24].

Finite element models (FEM) have been proposed to investigate the opto-mechanical interplay underlying CXL. The main challenge in developing accurate and reliable in silico models lies in the choice of the parameters that define the model’s mechanical response [25]. In the work of Roy and Dupps, the material parameters resulting from CXL-induced stiffening were determined by directly optimizing for the expected refractive outcome [26,27]. Wang and Chester published a multiphysics modeling of CXL [28] in which the mechanical properties were determined based on inflation tests performed on porcine eyes from the literature [29]. None of these models took into account the viscoelastic response of the tissue, nor was it calibrated on an independent set of data and subsequently applied to a clinical scenario.

In this study, we proposed a combination of mechanical testing techniques coupled with an inverse finite element modeling (iFEM) approach to determine the corneal anterior curvature changes induced by CXL. We combined a compression test (NI) with a high-resolution inflation test (OCE) to characterize the biomechanics of human cornea ex vivo before and after CXL, and we used this data to identify the parameters of a FEM aimed at replicating the biomechanical changes induced by the treatment. We then used the calibrated model to perform a patient-specific CXL simulation of advanced KC cases, taking into account not only the patients’ geometry but also the mechanical properties of the KC cornea, as determined with our in vivo OCE system [30]. The curvature changes predicted by the FEM simulations were compared with those measured at six-month follow-up to assess the applicability of the model in the clinical setting.

## Material and Methods

### 2.1 Ex vivo experiments

Corneas from five human donors (75±5.4 years, 60% female) were provided by the LIONS Cornea Bank Baden-Württemberg at the Eye Center, University of Freiburg. Donors or their legal representatives consented to its research use. The study was approved by the University of Freiburg’s ethics committee (408/15) and adhered to the principles of the Declaration of Helsinki. The corneas were deemed ineligible for transplantation and were stored in a cell culture medium for 45-65 days after collection. On the first day of testing, each cornea was subjected to NI and *ex vivo* OCE prior to undergoing CXL. To obtain an internal control, only half of each cornea was subjected to CXL treatment. On the second day of testing, NI and OCE were repeated on the CXL sample. The flowchart in Figure 1 summarizes the individual preparation and testing procedures.

**Figure 1.**
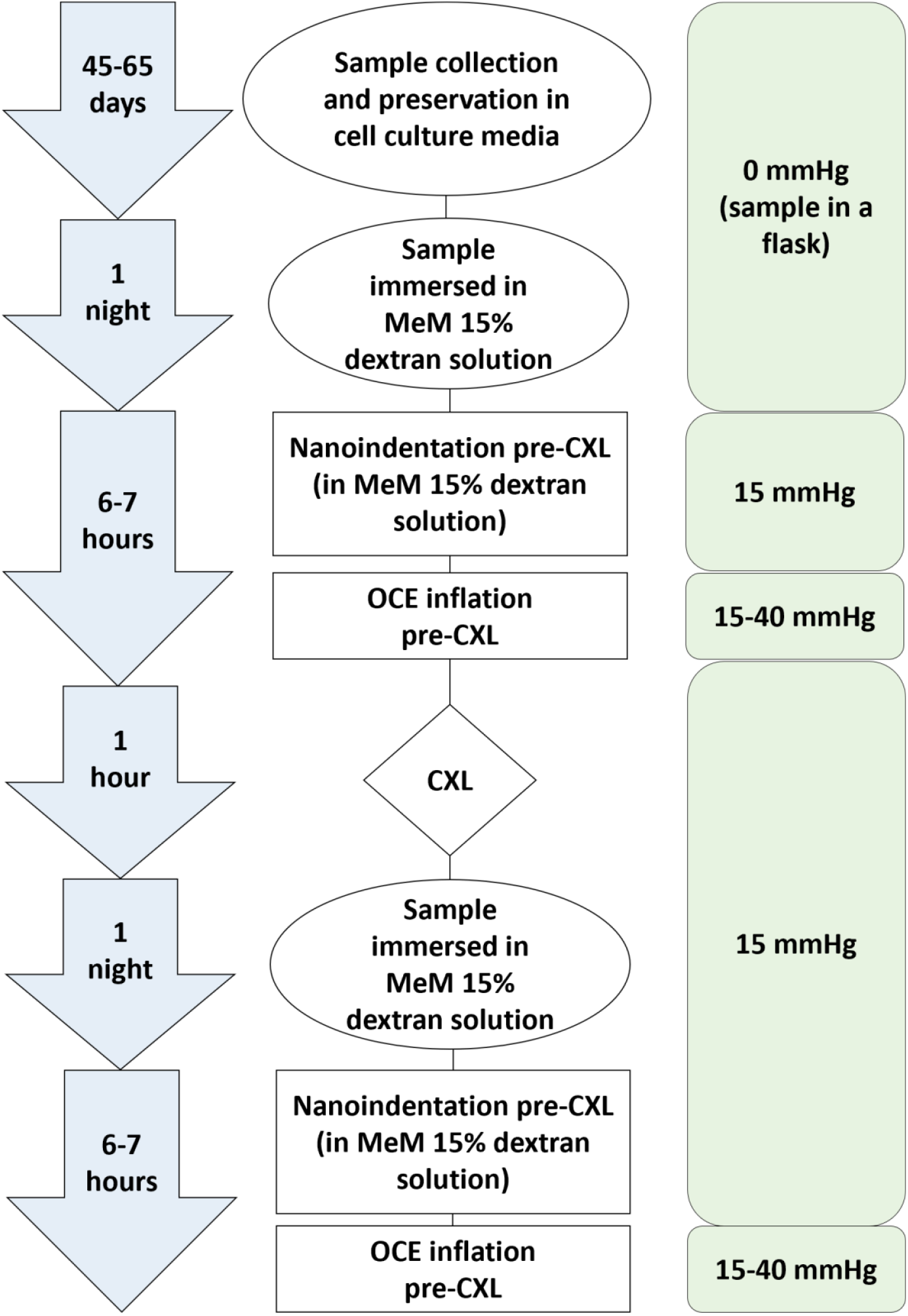
flowchart describing the experimental steps every sample was subjected to during the *ex vivo* experimental study phase. The duration of each step, the specific preservation solutions and the pressurization of the tissue are reported.

#### 2.1.1 Sample preparation

The night before the experiments, the sample was placed in a solution to restore physiological thickness. This solution consisted of minimal essential medium (MEM) earls cell culture medium (BS.F0325, Bio&SELL, GmbH, Feucht, Germany) supplemented with 15% Dextran 500 (9219.1, Carl Roth GmbH + Co. KG, Karlsruhe, Germany), 10% FBS SUPERIOR (FBS.S 0615, Bio&SELL, GmbH, Feucht, Germany), 100 U/mL penicillin and 100 µg/mL streptomycin (P4333-100ML, Sigma-Aldrich, St. Louis, MO, USA), 2,5 mg Amphotericin B (A2942-50ML, Sigma-Aldrich, St. Louis, MO, USA), 12,5 mM HEPES-Buffer (P05-01100, PAN-Biotech GmbH, Aidenbach, Germany) and 2 mM L-Glutamine (25030-081, Life Technologies, Paisley, UK). On the day of the experiment, the sample was mounted on a Barron K20-2125 artificial anterior chamber (Barron Precision Instruments L.L.C., Grand Blanc, MI, USA) and subjected to a retrocorneal pressure (RCP) of 15mmHg using a syringe connected to one side of the chamber. The other side of the chamber was connected to a pressure sensor (G19237 Mod. Duesseldorf, Geuder, Heidelberg, Germany). The corneal epithelium was removed with a blunt knife. The anterior chamber was placed on a custom-made sample holder that allowed the complete immersion of the sample in a MEM 15% dextran bath. The central corneal thickness (CCT) was measured at the beginning of each OCE experiment by optical coherence tomography (OCT) at a RCP of 15 mmHg.

#### 2.1.2 Nano-indentation

The cornea was subjected to NI testing (UNHT Bio – Anton Paar, Peseux, Switzerland), maintaining a RCP of 15 mmHg, with one side of the chamber connected to an infusion stand filled with saline. The NI was performed at 10 different sites in a 1 mm diameter area around the center of the cornea, with five sites on the irradiated half and five on the non-irradiated half of the cornea. The sample was fully immersed in the MEM 15% dextran solution during the NI experiment, which was performed at room temperature. The force of the indenter was controlled at a predetermined loading rate of 300 µN/min.

When the indenter depth reached a target threshold of 25 µm, the resulting maximum force was held for 90 seconds, then an unloading rate of 300 µN/min was applied. The force-displacement curves, the Hertzian modulus, *E*_*HZ*_ [Pa] and the indentation creep, *C*_*IT*_ [-] [22,31] were used to characterize the sample. A more detailed formulation of *E*_*HZ*_ and *C*_*IT*_ is provided in Appendix A.

#### 2.1.3 Optical coherence elastography

The anterior chamber was placed under an OCT system operating in the spectral range with a central wavelength of 877.8 nm, a bandwidth of 62.5 nm and an output power of 1.62 mW. The MEM 15% dextran solution was temporarily removed to avoid light distortion during the measurements. Starting from RCP of 15 mmHg, the cornea was inflated stepwise in 2 mmHg increments up to 40 mmHg. After each inflation step, a volumetric C-scan of a 12 × 12 mm area of the central cornea was acquired, consisting of a stack of 100 B-scans (axial resolution of 4.5 μm (in air), lateral resolution of 12.5 μm). Subsequent 3D tomographic images were compared to calculate the axial elastography strain *ε*_*zz*_ [-]. The phase-tracking algorithm [32–34] employed to obtain the elastography data has been described in previous publications by our group [20,35] and is described in Appendix B.

#### 2.1.4 Ex vivo CXL

A 0.1% riboflavin + 1.1% HPMC solution (MedioCROSS® M) was administered every two minutes for 30 minutes before the start of treatment to ensure sufficient uptake of the photosensitizer by the cornea. The solution was also administered continuously every two minutes during irradiation to prevent dehydration. A custom-made 1 cm-thick plastic mask covered half of the corneal surface to prevent exposure to UV light. The other half of the tissue was irradiated for 30 minutes with a 365 nm lamp (LED UV Curing System, Thorlabs, New Jersey) with an irradiance of 3 mW/cm^2^ (total fluence of 5.4 J/cm^2^).

### 2.2 Finite Element Model

#### 2.2.1 Material model

An incompressible, depth-varying, visco-hyperelastic material model was developed in FEBio (University of Utah, Weiss Biomechanics Lab and Columbia’s Musculoskeletal Biomechanics Laboratory, USA) [36] through an in-house built plugin [37]. The hyperelastic part of the model was described in a previous publication from our group [38] and detailed in Appendix C. Briefly, two dispersed collagen fiber families were included in the strain-energy function using a weighted integration of the Holzapfel-Gasser-Ogden (HGO) model [39] over the unit sphere [40] with 350 integration points:

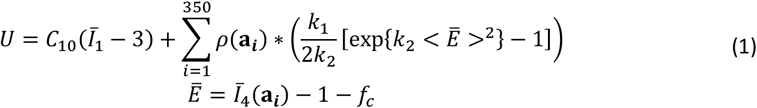

In this equation, the contribution of the extracellular matrix (first term of the equation) depends on the material parameter *C*_10_, while the second term of the equation describes the contribution of the collagen fibers to the strain-energy function and is based on the parameters *k*_1_, *k*_2_, and, *f*_*c*_ (fiber crimping). ***a*** corresponds to the general fiber direction in spherical coordinates. The angular density of the fiber distribution ρ(***a***) was expressed as the product of the in-plane and out-of-plane distributions *ρ*_*in*_ and *ρ*_*out*_. According to the findings of Nambiar and colleagues, the posterior cornea is 62% less stiff than the anterior one [41]. This biomechanical property of the tissue was enforced by linearly decreasing *k*_1_ across depth up to 38% of its anterior value 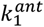.

The FEBio built-in solid viscoelastic model [42] was employed to describe cornea viscous behavior. According to this formulation, the second Piola Kirchhoff stress can be written as:

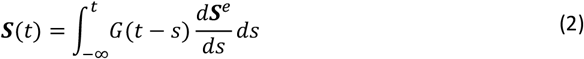

being ***S***^*e*^the (hyper)elastic stress and G(t) the relaxation function, expressed as:

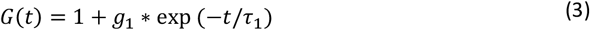

Where the coefficient *g*_1_ is the normalized viscoelastic coefficient. Therefore, the viscoelastic response of the material depends on the parameters *g*_1_ [-] and *τ*_1_ [s].

#### 2.2.2 CXL treatment model

The CXL treatment was modeled as a stiffening factor *K*_*cxl*_ multiplying the fiber-related parameter *k*_1_. To account for the depth dependence of the CXL treatment, *K*_*cxl*_ was linearly decreased across the corneal thickness until a value of 1 was reached at a depth of 300 µm. This position was chosen as an average value at which the demarcation line is observed in KC patients after the Dresden CXL treatment [43]. Not only was a depth dependence of the CXL effect modeled, but also a radial variation of the *K*_*cxl*_ factor, to mimic the radial attenuation of UV in the most peripheral part of the CXL area observed in the clinic. This variation results from the inclination of the corneal surface with respect to the irradiating beam. Following the approach described by Roy and Dupps, the parameter *K*_*cxl*_ was reduced with a Gaussian function and reached 10% of its value at a normalized radial distance of 0.9 from the center of the CXL zone [3].

#### 2.2.3 Parameter identification via iFEM

The material parameters *C*_10_, *k*_1_, *k*_2_ and *f*_*c*_ of the FE model were taken from previous works by our group in which iFEM was applied on UTT experimental data [38,41]. The fiber distribution parameters describing *ρ*_*in*_ and *ρ*_*out*_ were defined based on x-ray scattering and second harmonic generation data [44,45]. These parameters are summarized in Table 1.

**Table 1.** Material parameters adopted in the FEM for the pre- and post-CXL models.

| | $k$ | $C_{10}$ | $k_1^{ant}$ | $k_2$ | $f_c$ | $a_{ant}$ | $a_{pos}$ | $b$ | $g_1$ | $\tau_1$ | $K_{CXL}$ |
| --- | --- | --- | --- | --- | --- | --- | --- | --- | --- | --- | --- |
|  | [MPa] | [kPa] | [MPa] | [-] | [-] | [-] | [-] | [-] | [-] | [s] | [-] |
| Pre-CXL | $10^3$ | 25 | 4.56 | 18.9 | 0.2 | 0 | 5 | 2.5 | 3.89 | 29.32 | 1 |
| Post-CXL | $10^3$ | 25 | 4.56 | 18.9 | 0.2 | 0 | 5 | 2.5 | 6.83 | 10.18 | 16.3 |

To determine the viscoelastic parameters *g*_1_ and *τ*_1_, the creep NI experiments of the pre-CXL samples were modeled in FEBio. An average emmetropic 3D corneal geometry was defined [46] based on two spheres cut with theta of 50° representing the anterior and posterior surface (anterior radius of 7.5 mm, posterior radius of 6.4 mm, CCT of 500 µm), together with a rigid spherical indenter of 0.5 mm radius (Appendix D). The geometries were meshed with 7208 and 2048 hexahedral linear elements (HEX8) for the cornea and indenter, respectively. The indenter retained only its vertical translational degree of freedom and was fitted with the average force profile experimentally imposed on the indenter. To mimic the condition of the sample mounted on the anterior chamber, the peripheral junctions corresponding to the cornea-limbus were constrained by applying zero-displacement and zero-rotation boundary conditions along each axis, and the posterior surface was inflated to 15 mmHg.

We minimized the Mean Square Error (MSE) between the average experimental data and the displacement of the indenter calculated using FEM with a Nelder Mead optimization algorithm using the Scipy built-in optimization library [47]. The unloading phase was not considered in the optimization due to the artifacts caused by dextran capillary force acting on the indenter and the viscosity of the tissue itself. A total of three optimizations were run: i) the parameters *g*_1_ and *τ*_1_ were determined considering the NI experiments with the cornea in its untreated state. ii) The entire corneal model was then cross-linked and a second optimization was carried out using the experimental post-CXL-NI data to determine the stiffening factor *K*_*CXL*_. iii) A third optimization was then performed, keeping the optimized *K*_*CXL*_ factor and leaving *g*_1_ and *τ*_1_ free to determine whether the viscous parameters were affected by CXL.

After the model parameters were determined using iFEM, the OCE inflation test was simulated on the average corneal model to validate the FEM. The peripheral nodes corresponding to the corneal-limbal junction were constrained by applying zero-displacement and zero-rotation boundary conditions along each axis, and the posterior surface was inflated to 15 mmHg. Inflation steps of 2 mmHg were then applied to the posterior surface, mimicking the OCE measurements. The axial strains obtained from the OCE experiments, averaged over the anterior 300 µm of the tissue, were compared to the FEM-derived strains in the z-direction, averaged over the same region of the model, for both pre- and post-CXL treatment data.

### 2.3 Clinical validation of the model

Three patients with advanced KC (Table 2) who were eligible for CXL treatment were considered for this study, which was approved by the Cantonal Ethics Committee of the Canton of Zurich (January 11, 2022, number 2021-02275). Informed consent was obtained for each subject and each dataset was pseudonymized prior to analysis. Each subject underwent an *in vivo* OCE assessment before receiving individualized CXL treatment on the same day. Corneal topographies (Pentacam, OCULUS Optikgeräte GmbH) were taken one hour before treatment and at the six-month follow-up visit.

**Table 2.**
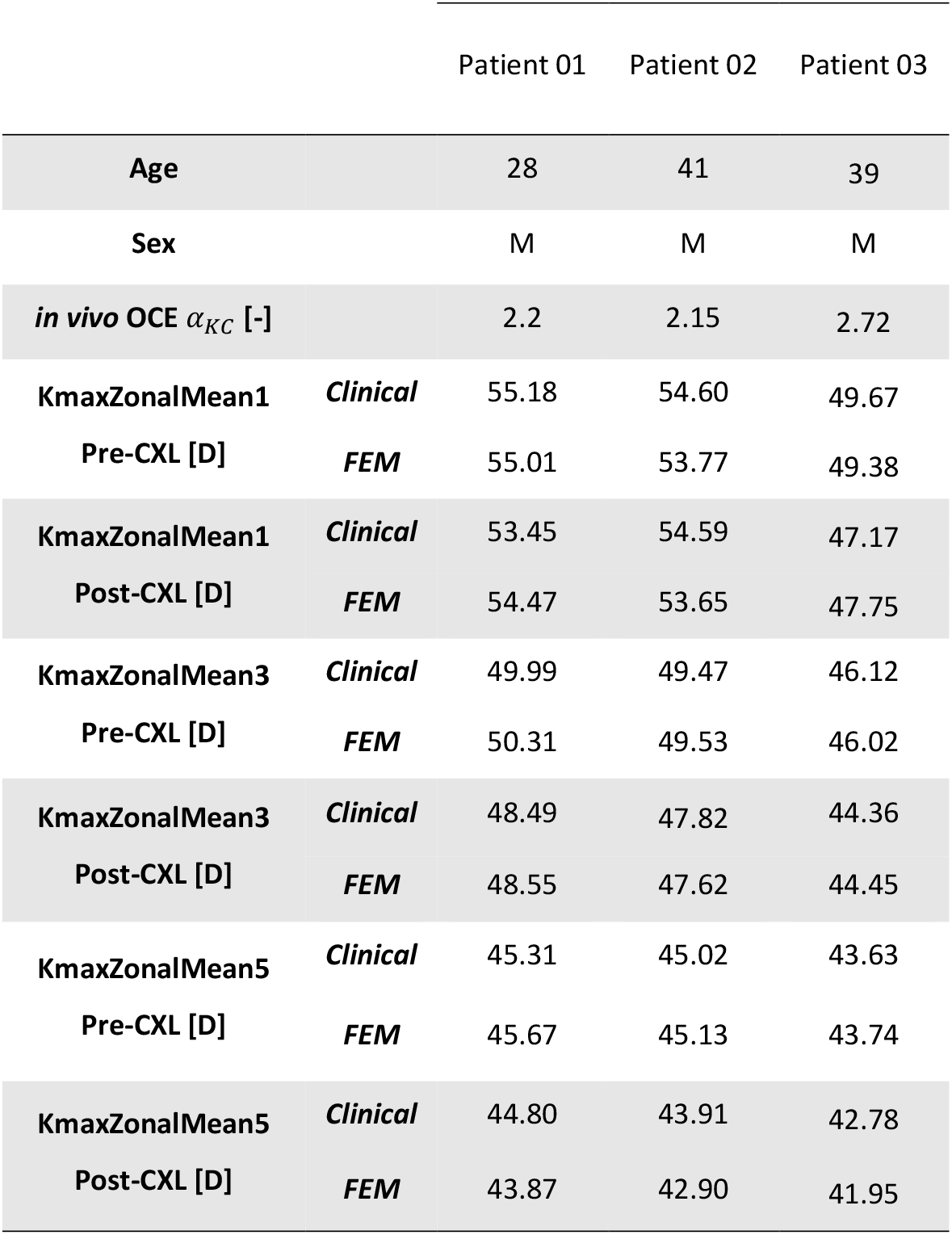
Demographic, clinical and FE-derived data of the three patients analysed in this study.

#### 2.3.1 In vivo OCE

The *in vivo* OCE measurements were acquired with a setup previously described [30,48], which differs in the OCT scan employed and measurement method w.r.t the *ex vivo* OCE setup described in paragraph 2.1.3. In brief, the patient sat in front of a commercial spectral-domain anterior segment OCT machine (ANTERION, Heidelberg Engineering, Germany) operating at a central wavelength of *λ*_*mean*_=1300 nm and provided with an axial and lateral resolution of 9.5 µm (in air) and 30 µm, respectively. Before the OCT scan, the patient was asked to wear custom-made polycarbonate swimming goggles (n of 1.586) and to look at the internal fixation light (Maltese cross) of the device during the entire examination period. After the system was started by the operator, 128 consecutive OCT B-scans were acquired along a line running through the center of the cornea (A-line rate of 50 kHz). After approximately 1/3 of the examination, the system was automatically triggered to immediately cause a slight dilation of the eye by lowering the ambient pressure in the eyeglass chamber by 10 mmHg.

Post-processing of the dispersion-corrected complex-valued OCT scans was performed by comparing consecutive B-scans using the same phase-tracking approach described in paragraph 2.1.3 and Appendix B to obtain the *in vivo* derived strain map *ε*_*zz*_ (*z, x*). Once the axial strain distribution was obtained, we calculated the ratio *α*_*KC*_ [-] between the strain within the cone region 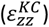 and the rest of the cornea 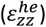:

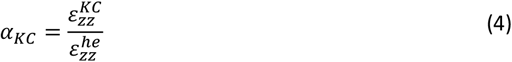

The ratio *α*_*KC*_ represents the index of mechanical weakening. If *α*_*KC*_ >0, the KC region deforms more than the rest of the cornea when exposed to the same Δp.

#### 2.3.2 Second-generation Customized CXL protocol

Patients received the ELZA-PACE CXL protocol, a procedure that employs gradients of riboflavin and UV light to achieve corneal flattening and asymmetry reduction. This technique creates multiple gradients to achieve a tailored effect: in summary, the PTK region serves as an epi-off CXL area, contrasting with the rest of the cornea, which is treated using an epi-on approach. First, an excimer laser is used to perform an epithelial map-driven PTK, which selectively removes a limited area of epithelium over the cone while preserving the underlying stromal tissue, effectively creating a partial epi-on/epi-off procedure. The epithelial map was generated from images obtained using the anterior segment optical coherence tomographer combined with a Placido disc (MS 39, CSO Italy, Italy). Next, a riboflavin soaking was performed using hypo-osmolaric riboflavin 0.1% (Riboker, EMAGine AG, Switzerland) and a riboflavin concentration gradient was established, with the highest levels concentrated directly over the cone. Finally, the tissue was irradiated with UVA light with a C-eye device (EMAGine, Switzerland) that was guided by topography and epithelial thickness maps to irradiate a 4 mm diameter spot centered on the KC cone with a fluence of 15 J/cm^2^ and a larger 9 mm diameter area at 8.1 J/cm^2^.

#### 2.3.3 Patient-specific finite element modeling

The finite element mesh was developed in Python using the Gmsh library [49] and was based on the preoperative cornea topographies (Appendix D). The total number of second-order tetrahedral elements (10 nodes per element) was approximately 150000, and the geometries were constructed such that the mesh was finer in the central region with a 3 mm radius area while becoming coarser towards the periphery. The number of elements across the central corneal thickness was seven. Sliding boundary conditions were applied to the nodes corresponding to the limbus region by only allowing radial displacements in a spherical coordinate system [50]. The KC region was identified on the anterior and posterior surfaces by the nodes whose tangential curvature exceeded the 67.5^th^ percentile of the curvature distribution, considering only a zone of 8 mm diameter around the center of the cornea. After identifying the two nodal regions corresponding to the apex and base of the cone, a convex hull was created using these regions as boundaries. The elements whose centroids lay within this convex hull were then assigned a KC flag. The material model described in subsection 2.2.1 was applied, dividing the *k*_1_ parameter in the KC region by the corresponding reduction in axial strain *α*_*KC*_ measured with the *in vivo* OCE. The change of *k*_1_ was implemented gradually, assigning the *in vivo* reduction to the center of the KC region and then linearly decreasing this value to obtain no reduction at the transition zone.

The model was then applied a pre-stress algorithm to retrieve the initial (stress-free) geometry required to achieve the patient-specific geometry after pressurizing the posterior surface at 15 mmHg (normal IOP) [51]. After obtaining the initial configuration, CXL was applied to the model and normal IOP loading (15 mmHg) was simulated. The treatment was centered on the cone region and two different concentric circles were defined, mimicking the ELZA-PACE-CXL protocol described in paragraph 2.3.2. To account for the high fluence delivered in the innermost circle, the CXL depth in this region was increased to 400 µm, which corresponds to the demarcation line position within the KC area for patients undergoing ELZA-PACE-CXL. In addition, the *K*_*cxl*_ factor, which was experimentally calibrated based on the Dresden CXL, was increased by 33% within the inner region of 4 mm diameter. This adjustment was derived from the results of UTT on full-thickness porcine specimens. These tests showed a 33% increase in the stiffening effect after CXL at a fluence of 15 J/cm^2^ compared to the standard Dresden protocol when adjusted for the depth of the cornea that was actually crosslinked [52]. As the simulation was quasi-static, the viscous part of the model was excluded.

The FEM-derived CXL-induced geometric changes were compared with the clinical ones at the six-month follow-up. To quantify CXL-related topographic changes, we adopted KmaxZonalMean1, KmaxZonalMean3, and KmaxZonalMean5 [D], which represent the average corneal curvature within a 1, 3, and 5 mm diameter regions centered around the point of maximum curvature, respectively [53]. Changes in the anterior corneal KmaxZonalMean and in the tangential curvature *K*_*tg*_ [D] map, both derived from the simulation, were compared with the corresponding values obtained from the topography data at the six-month follow-up.

### 2.4 Statistical analysis

The statistical analyses were performed in Python. Continuous variables were expressed as means (± standard deviation) or medians (Q1-Q3). Paired Wilcoxon tests were used to compare stiffness and creep measures (*E*_*HZ*_, *ε*_*zz*_, *C*_*IT*_) before and after CXL for the UVA-irradiated region. A p-value of <0.05 (α) was considered statistically significant. Figures were generated in Python, Matlab, and FEBio Studio.

## Results

### 3.1 Ex vivo experiments

The CCT was 419±60 µm and 484±33 µm (p=0.125) before and after CXL, respectively. When comparing the mechanical behavior in the CXL region of the cornea, the NI experiments showed an increase of 41.0±16.6 kPa in *E*_*HZ*_, with values increasing from 99.96±15.8 kPa to 141.0± 4.7 kPa (p<0.01) before and after CXL, respectively, as shown in Figure 2A. *C*_*IT*_ decreased by 2.5±1.7 % after CXL, from 19.02±2.0 % in the untreated cornea to 16.48±1.3 % (p<0.01) after treatment, as shown in Figure 2B. Of the 25 indentation sites considered, one was excluded from the analysis due to incorrect contact detection by the indenter.

**Figure 2.**
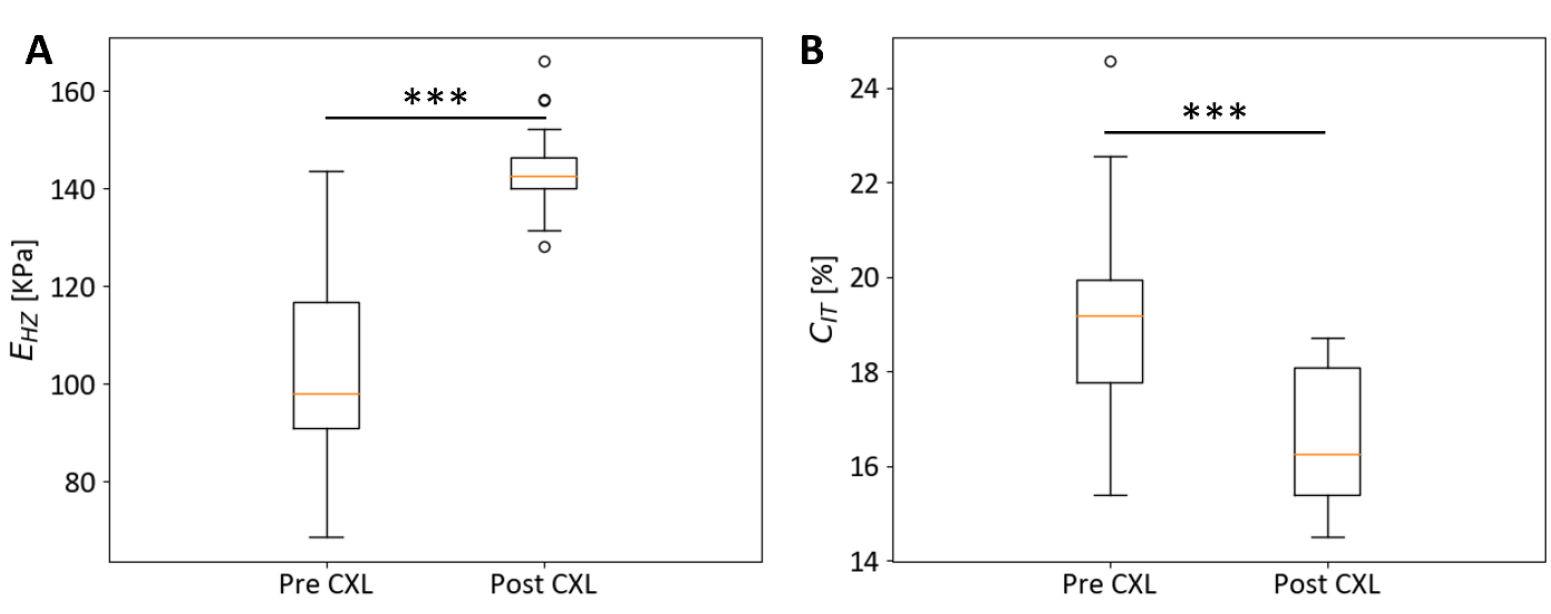
NI experimental results collected on n=24 data points (5 samples and 5 measurements per sample, one measurement excluded). A) Elastic Hertz modulus *E*_*HZ*_. B) indentation creep *C*_*IT*_. ***.001≤p≤.01.

In the UVA-irradiated region of the cornea, the high-resolution OCE showed reduced strain amplitudes in the first 300 µm in post-CXL tissue when compared to pre-CXL condition (−5.92±0.8‰ vs 1.1±2.6‰; p=0.06, respectively).

### 3.2 Mechanical parameter identification

Before CXL, the iFEM model closely matched NI experimental loading and holding curves, staying within one standard deviation of experimental values except at low displacements (<7 µm), where it slightly overestimated. After CXL, the model underestimated experimental values at lower strains but matched the experiments in the holding phase (Figure 3A). The optimized parameters are listed in Table 1. The three iFEM optimizations required 207 (MSE=0.016), 65 (MSE=0.12), and 202 (MSE=0.059) iterations to retrieve *g*_1_ and *τ*_1_ in the native condition, the stiffening factor *K*_*CXL*_, and *g*_1_ and *τ*_1_ after the treatment, respectively. The CXL resulted in a 16-fold stiffening of the fiber-related modeling parameter *k*_1_, as well as an increase in *g*_1_ (+175%) and a decrease in *τ*_1_ (−288%).

**Figure 3.**
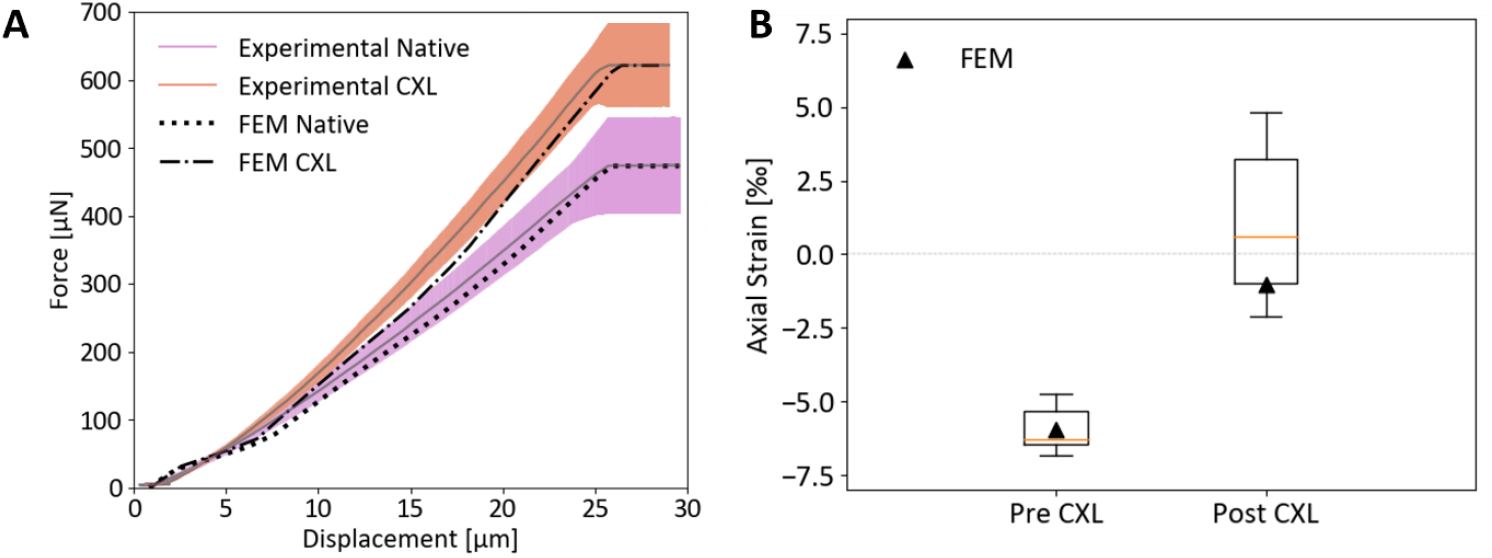
Comparison between experimental and numerical data collected on n=5 samples. A) NI force-displacement curves used to tune the model. Dashed and dotted lines represent the numerical response. B) OCE axial strain used for model validation (boxplots) compared to the strain calculated with the FE model (black triangle marker).

The calibrated model was adopted to simulate the OCE inflation test. The variation in axial strain induced by the CXL model (−5.94‰ vs. −1.0‰ before and after CXL) is shown in Figure 3B. The numerical results fell within the range of the *ex vivo* OCE data, with the native condition nearly matching the mean experimental value, while the post-CXL condition reached the 25^th^ percentile of the distribution. The *ε*_*zz*_ cross-sectional distributions as obtained from the *ex vivo* OCE are compared with those determined by FEM, either before and after CXL (Figure 4). When inflated from 15 to 17 mmHg, the cornea experienced an axial compression across its entire thickness. This behavior was fully captured by the model in the native state. The CXL OCE cross-sectional view shows how the treatment affected only the anterior 300 µm of the cornea. Compared to the experimental data, the CXL model showed a softer response to the inflation, with less axial strain in both the non-CXL and CXL regions.

**Figure 4.**
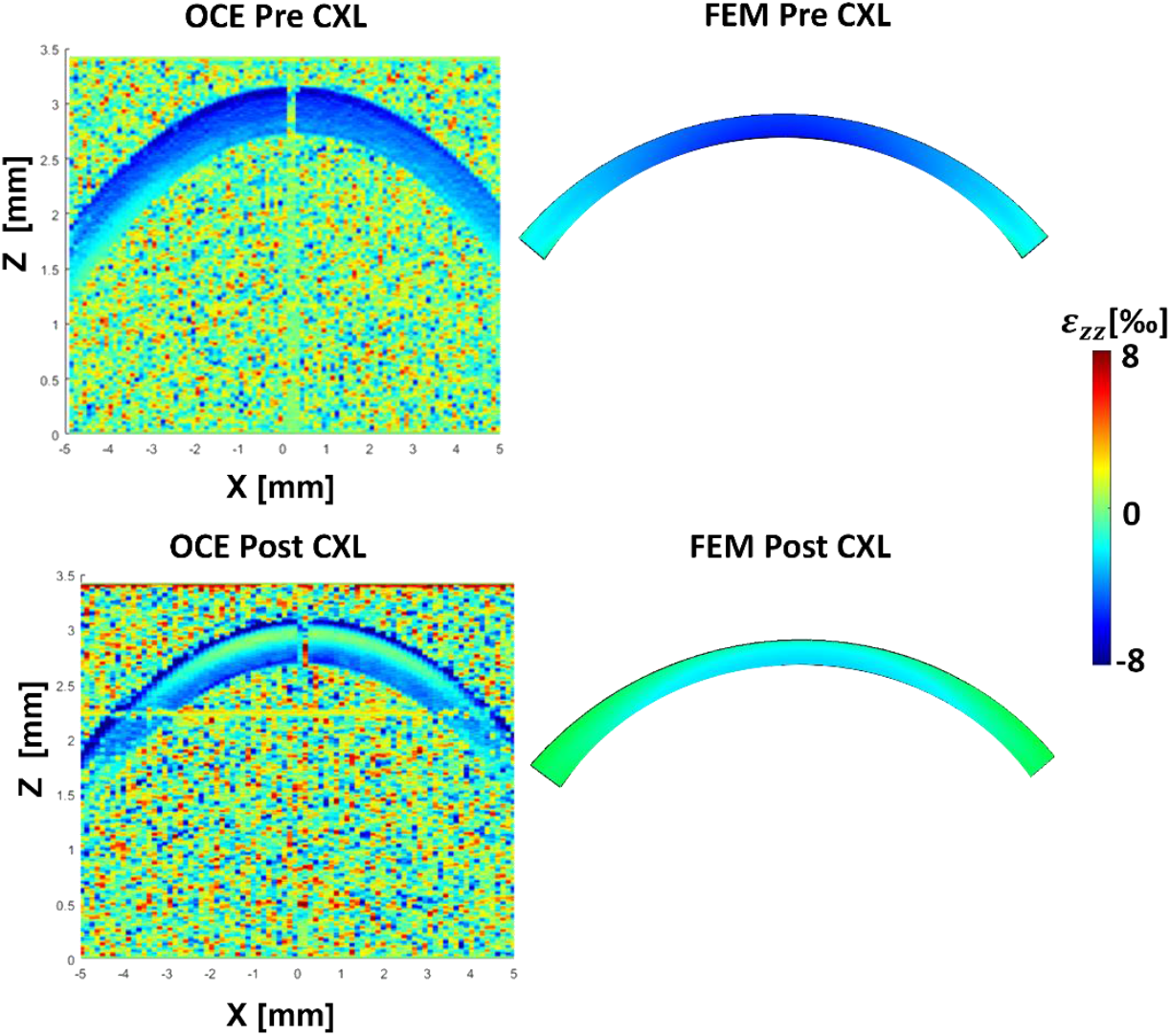
Cross-sectional map of the OCE strain of a representative sample (left column) compared to FE results (right column), calculated with a pressure gradient of 2 mmHg. The first row shows the state before CXL, the second row the state after CXL. The color scale shows the axial strain *ε*_*zz*_ [‰] values.

### 3.3 Clinical validation of the model

The result of the *in vivo* OCE analysis is shown for the three patients, represented by a B-scan colored with the *ε*_*zz*_ distribution due to the induced cornea inflation (Figure 5). The ratio *α*_*KC*_ between the axial strain of the cone region and the rest of the cornea is listed in Table 2 for the three patients. On average, we recorded a 2.36 times higher deformation in the cone region when the patient was exposed to Δp=10 mmHg, with comparable values recorded for the different patients, ranging from 2.15 to 2.72. When applied to the model by dividing *k*_1_ for the ratio *α*_*KC*_, this resulted in a reduction of *k*_1_ up to 45.4%, 46.5%, and 36.7% of its original value for patients 01, 02, and 03, respectively.

**Figure 5.**
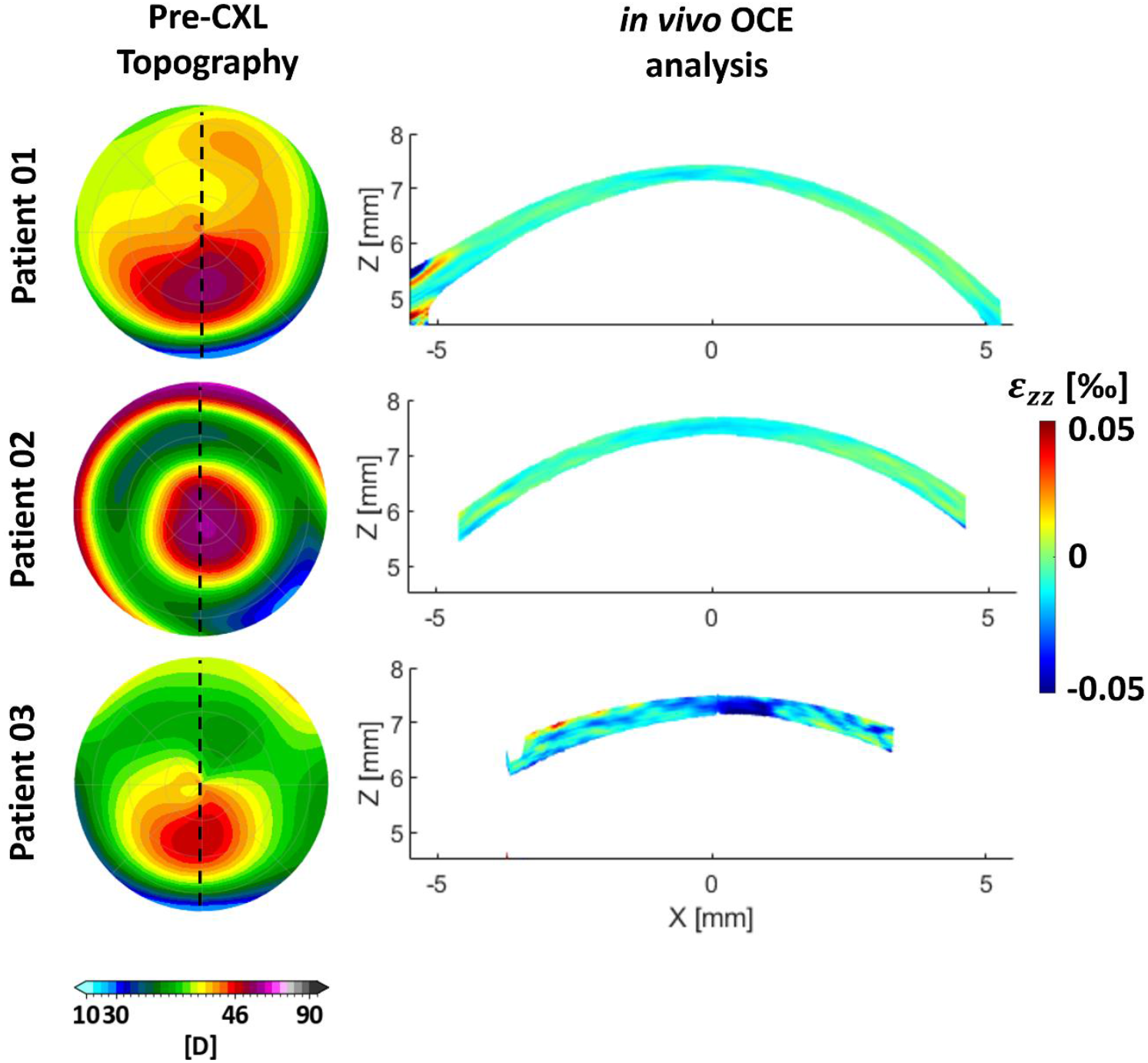
*in vivo* OCE analysis results. Left column: enface view of the pre-surgical anterior cornea curvature. The black dotted line represents the position to which the cross-sectional scan was taken. Right column: *in vivo* OCE axial strain maps. Color scale represents axial strain *ε*_*zz*_ [‰] values.

The model accurately reproduced the 6-month follow-up clinical data, especially for patient 02, as shown in the tangential curvature maps in Figure 6, particularly in the region of the cornea surrounding the cone, while differences in the order of 1 or 2 D are reported in peripheral areas, where the meshes are coarser. The comparison between clinical and FEM ΔKmaxZonalMean3 for the three patients was −1.5 D vs −1.76 D, −1.65 D vs −1.91 D and −1.76 D vs −1.57 D (Table 2).

**Figure 6.**
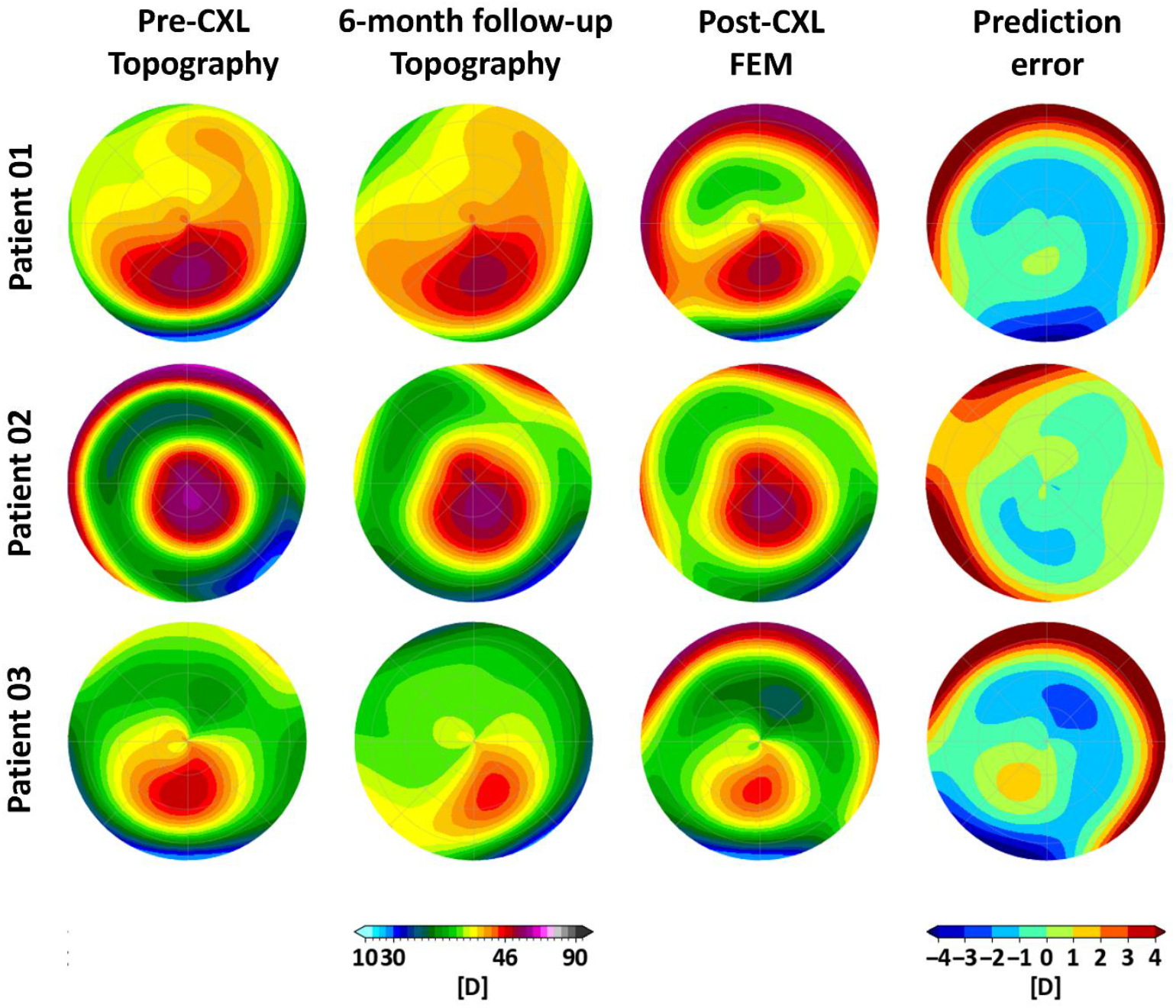
Anterior surface tangential curvature maps [D] obtained on a 7 mm diameter optical zone for each of the three patients (one per row). Left colum represents the Pentacam topography before CXL, central left column the Pentacam topography at 6-month follow-up, central right column the post-CXL FEM result, right column the difference between the FEM-predicted anterior curvature and the clinical curvature at follow-up.

## Discussion

CXL-induced refractive changes are not taken into account during preoperative planning in the clinical setting, making it difficult to accurately predict optical performance after treatment. In this context, FEM could serve as an effective prediction solution to bridge this gap. We propose a patient-specific FEM that captures the mechanical effects induced by CXL. A strength of this model is that it has been calibrated and validated against various mechanical test methods that account for both tensile and compressive tissue behavior. In addition, a patient-specific *in vivo* mechanical analysis was included to tailor the mechanical properties to the individual patient. Consequently, the proposed FEM is customized in both shape and mechanical properties, paving the way for a predictive tool that truly reflects the individual patient’s condition.

NI is a well-known technique for investigating the viscoelastic properties of the cornea. Nohava et al. reported the CXL-induced corneal changes using a similar NI instrumentation and showed a 15 kPa increase in elastic modulus and a 10% decrease in creep after procedure [22]. These data differ from the 41 kPa increase and 2.5% decrease of *E*_*HZ*_ and *C*_*IT*_, respectively, reported in the present study. A common disadvantage of performing NI is the fixation of the sample on a Petri dish, which does not correspond to the physiological, pre-stretched state of the tissue. While Nohava et al. glued their samples to a Petri dish, our setup allowed testing the cornea while mounted on the anterior chamber and applying a specific RCP. In the experiments described in the present work, the tissue was pre-stretched under a physiology-like load of 15 mmHg, which can explain the difference in CXL effect between these two studies.

The stiffness of the fiber-related component in the HGO model increased by a factor of 16 after CXL, resulting in a 105% increase in tangential modulus at 15 mmHg on the apical rise curve. This value is lower than the 200-300% increase in elastic modulus reported by Roy and Dupps after an inverse analysis on 16 patients with KC [27]. In another computational study, the same authors showed that a 200-300% increase in Young modulus was necessary to achieve a 2 to 3 D flattening of the steepest corneal point [3]. In the present study the achieved tangential curvature reduction was lower (−1.75 D on average), which is in agreement with the lower increase in stiffness assigned in our CXL model. Wollensak et al. reported a 350% increase in Young modulus at 6% strain after CXL on human corneas measured by UTT [21]. The stiffness increase reported in the present study was calculated at a stress of 2 kPa (equivalent to 15 mmHg), which is much lower than the stresses induced by Wollensak et al. using UTT, which probably explains the lower CXL effect of the present model due to the nonlinear stress-strain behavior of the corneal tissue. Kling et al. reported a 36.86% increase in Young’s modulus on porcine corneas after CXL, using a simple biomechanical model and inflation tests [29]. However, due to the different material models and the use of porcine tissue, direct comparison with our findings is not feasible. In summary, significant variations in the stiffening effect of CXL have been reported in the literature due to different protocols and sample types. For this reason, we performed experimental and numerical studies tailored to our clinical protocol. Despite the use of a single stiffness value, our experimental data show that the interindividual variation is much smaller than the effect of CXL, and our numerical model successfully replicates the experimental behavior.

The experimentally determined reduction in creep behaviour was reflected in the model by a threefold reduction in the parameter *τ*_1_ after CXL. In NI experiments, not only the fabric relaxation was measured, but also the poroelastic component caused by the fluid flow under the indentation force [54]. However, the corresponding viscoelastic FEM parameters did not distinguish between the viscoelastic and poroelastic responses of the cornea, as the adopted viscoelastic model is not related to specific material parameters such as viscosity or hydraulic permeability. Therefore, our model is phenomenological, which is a limitation of this approach.

The reduction in axial strain magnitude assessed by OCE confirms previous findings by our group in rabbit and porcine eyes [20,55]. An important result of this study is the comparison between *ex vivo* OCE and FEM, which served as validation for the iFEM algorithm. These data not only demonstrated the accuracy of the model, but also proved the validity of OCE in quantitatively assessing tissue mechanics with high-resolution at low deformations.

Compared to previous CXL computational models [26,27], the present FEM takes into account the viscoelastic behavior of the cornea before and after treatment and was adapted and validated using various experimental tests on human specimens. In addition, the depth dependence of the corneal stiffness was modeled. After calibrating the model, we proposed its clinical application by implementing a Python-driven pipeline to perform patient-specific CXL simulations. This tool is fully automated and relies entirely on open-source software for meshing, pre- and post-processing and finite element analysis. The entire process takes about three hours and can be run on any operating system. We tailored the simulation to the individual patient by replicating the corneal geometry from clinical topographies and performing an *in vivo* mechanical analysis to tune the KC material properties. To our knowledge, this is the first time that *in vivo* preoperative OCE data has been used as input to a FEM to predict topographic outcome and compare it to clinical postoperative data. The KC region was measured to be 57.1% softer than the rest of the cornea in terms of axial strain amplitude, which is consistent with previous reports: UTT [7] and acoustic radiation force elasticity microscopy [11] were used to measure KC buttons that were ∼50-70% softer than the healthy cornea. Similar attenuation factors were reported by Scarcelli et al. who observed ∼70% reduction in shear modulus measured by Brillouin microscopy both *ex vivo* and *in vivo* [9,10]. The *in vivo* OCE data from patient 03 were noisier than the others, most likely due to the greater deformation of this cornea under pressure. Even though we only report on a small cohort of three patients, our results prove that OCE is a valid tool for *in vivo* biomechanical assessment of corneal stiffness. Combining this analysis with a polarization-sensitive OCT setup [56] could allow for the measurement of patient-specific fiber distributions, which are absent in the present work, further tailoring the model to the single patient’s condition.

We assumed that the complex genetic, biochemical and ultrastructural mechanisms underlying KC can be fully modeled as local stiffness reduction [3]. This approach is suitable since the aim of this model is not to represent the progression/regression of KC over time, but to evaluate the effects of CXL treatment at a specific time point. Tangential curvature was used to detect and delineate the KC area, as it has been shown to be a more reliable index than sagittal curvature in KC detection [57,58]. As a consequence of mechanical stiffening, localized CXL has been shown to flatten the cone and improve visual acuity [59,60]. Our model was able to replicate the dioptric correction induced by individual CXL by reproducing the clinical stiffening gradient in the cornea. The correction predicted by the FEM appears to slightly overestimate the clinical correction at the six-month follow-up, but a larger number of patients need to be studied to verify this tendency.

The experimental CXL was limited to half of the cornea. This allowed the cutting of strips in both the irradiated and non-irradiated areas of the tissue, which were tested with UTT. Appendix E details the methods and results of the UTT experiments. Surprisingly, no statistically significant differences were found between the irradiated and non-irradiated areas of the cornea after CXL, neither with UTT, NI nor OCE, as shown in Figures E1-4. Most likely, it was the case that the in-house manufactured mask to prevent the light from irradiating half of the cornea failed to properly shield the tissue, allowing UVA light to be scattered, which could explain why the non-irradiated half of the cornea became stiffer and therefore underwent some amount of CXL. In the planning phase of the study, the UTT data should be included in the iFEM. These data were latterly removed from the optimization as it was not possible to exclude the presence of CXL in the “non-treated” region. The data and comparisons reported in the present study are therefore limited to the comparison between pre- and post-treatment in the irradiated part of the cornea.

Despite the promising results, the present study is not free of limitations. The FEM used for the iFEM was built on an average human corneal geometry, with no zero-load geometry derivation, and the algorithm was run on the average experimental responses rather than on each individual *ex vivo* sample. This averaging approach may have hindered geometric differences between the analyzed samples. We also reported an increase in CCT, although not statistically significant, before and after CXL. This geometric feature was not included in the average model. Therefore, the iFEM result only reported the changes due to the CXL effect, while the thickening, which in reality contributes to the stiffening experimentally determined via NI and OCE, is not considered. Moreover, the Dextran solution, as well as the preservation condition, might have had an influence on the mechanical properties of the *ex vivo* samples by affecting their pre-CXL thickness and hydration levels.

Another limitation was that the experimental CXL data used to calibrate the model came from healthy donor corneas, which were individuals significantly older than the KC patients who participated in the study. The model parameters were identified using corneas that were naturally stiffer than those of the KC patients, due to both the advanced age of the donors and the lack of biomechanical weakening caused by the KC disease itself. Consequently, the model may not have fully captured the mechanical effect of CXL on the less stiff and more compliant corneas typically seen in younger KC patients. Similarly, the model’s stiffening factor was calibrated to the Dresden CXL performed *ex vivo*. When scaled to the high-energy customized ELZA-PACE-CXL, this factor was increased by 33% based on previous experiments conducted on CXL protocols with comparable fluences [52]. However, these tests were conducted *ex vivo* on porcine corneas, and therefore may not be representative of the actual stiffening difference between the two clinical protocols. These approximations may explain the imperfect agreement between model predictions and clinical results, and diminish the patient-specific relevance of the model. More *ex vivo* experiments are required to better characterize different CXL protocols, but the characterization of KC corneal samples is very difficult due to the challenges associated with obtaining such tissue for research purposes. For this reason, the *in vivo* quantification proposed in this study is attractive despite its technical difficulties.

The *in vivo* quantification of CXL-induced curvature changes was conducted on three patients, making statistical analysis unfeasible. Additionally, IOP measurements were unavailable for these patients, leading to the use of a standard value of 15 mmHg, which may have hindered inter-individual loading variations. While this did not impact the *in vivo* OCE measurements, which assess the relative deformation between the two tissue regions, it did affect the FEM, which considered the same loading condition for each individual. Furthermore, the present study lacked *in vivo* OCE post-CXL data, which could have been employed to evaluate the treatment efficacy. Given the unavailability of data on the procedure’s success, each CXL treatment was assumed clinically effective and modeled solely as a mechanical stiffening of the cornea, without accounting for the biological remodeling processes commonly associated with this procedure.

Since the three optimizations on an HPC cluster with 16 CPUs and 32 GB RAM took 47.9 h, 13.5 h and 46 h, respectively, a major drawback of the iFEM approach is its time inefficiency, especially when optimizing for more than one parameter. Physics-informed neural networks have shown promise in accelerating inverse problem solutions [61] and could potentially help address this limitation in the future.

## Conclusions

A novel patient-specific CXL model was developed. It was calibrated and validated by combining compression and inflation tests to fully characterize the time-dependent and nonlinear mechanical behavior of the cornea. The model was applied in a clinical setting by tailoring the geometries and mechanical properties measured *in vivo* to the specific patients. The close agreement between the topographic correction achieved by CXL treatment six month after treatment and the prediction of the model shows that the proposed FEM can improve the planning of customized CXL procedures.

## Declarations

### Ethics approval and consent to participate

Ethical approval for both studies described in this manuscript was obtained as detailed in the corresponding Methods sections. All procedures were conducted in accordance with the relevant institutional and national ethical standards. Clinical trial number: not applicable.

### Consent for publication

All authors have given their informed consent for publication of this manuscript.

### Availability of data and material

The raw data supporting the conclusions of this article are available from the authors upon reasonable request, without undue restrictions.

### Competing interests

None declared. The authors declare that they have no known competing financial interests or personal relationships that could have appeared to influence the work reported in this paper.

### Funding

This work received funding from the European Union’s HORIZON 2020 research and innovation programme under grant agreement No 956720. Additional funding was provided by Cusanuswerk Bischöfliche Studienförderung through a PhD scholarship and travel grant

### Authors’ contributions

MF: Conceptualization, Data curation, Investigation, Methodology, Software, Validation, Writing– original draft, Writing–review and editing.

RL: Data curation, Investigation, Methodology, Software, Validation, Writing–review and editing.

MA: Data curation, Methodology, Software, Writing–review and editing.

GS:: Methodology, Investigation, Supervision, Writing–review and editing. SL: Methodology, Investigation, Supervision, Writing–review and editing.

ET: Clinical data collection, Methodology, Investigation, Supervision, Writing–review and editing.

FH: Clinical data collection, Methodology, Investigation, Supervision, Writing–review and editing.

PB: Conceptualization, Investigation, Methodology, Project administration, Resources, Supervision, Validation, Writing–review and editing.

SK: Conceptualization, Formal Analysis, Funding acquisition, Investigation, Methodology, Project administration, Resources, Software, Supervision, Validation, Writing–review and editing, Writing– original draft.

## Declaration of generative AI and AI-assisted technologies in the writing process

During the preparation of this work the authors used ChatGPT-4o in order to improve language and readability. After using this tool, the authors reviewed and edited the content as needed and take full responsibility for the content of the publication.

## Supplementary Material

## Appendix A

In the NI experiments, the Hertz’s modulus *E*_*HZ*_ was obtained by fitting the following equation to the force-depth curve in the region encompassed between 10% and 98% of the maximum load:

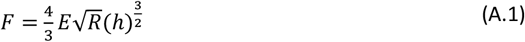

*F* rapresents the indentation force, *R* the radius of the indenter tip (0.5 mm), and *h* the indentation depth.

The indentation creep *C*_*IT*_ was calculated according to the standard ISO 14577:

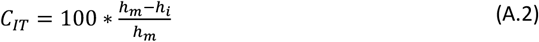

where the indentation depths at the beginning and at the end of the hold period and given by *h*_*i*_ and *h*_*m*_, respectively.

## Appendix B

The algorithm for the OCE strain computation is detailed herein. Briefly, a phase-sensitive deformation tracking algorithm was used to obtain the phase difference between the corresponding A-scans in the reference and deformed (Δp=2 mmHg) configurations by amplitude-weighted complex cross-correlation:

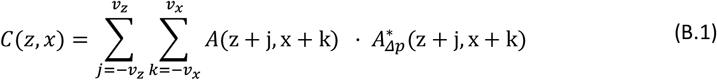

Where A(z, x) is the complex OCT interference signal acquired at position (z,x) [m] and v_z_ = 3 and v_x_ = 3 [pixels] is the size of the phase-processing windows used. The phase difference was employed to obtain the pixel-wise strain in the direction of the optical axis, *ε*_*zz*_ [-] by deriving the axial displacement U(z,x) [m] along z:

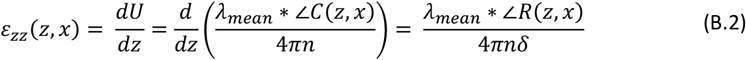

Where n = 1.375 [-] is the refractive index of the cornea, δ = 4.48 μm is the OCT axial sampling unit (in air), ∠R is the angle of a second complex cross-correlation 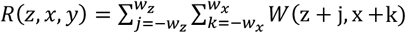 and w_z_ = 3 and w_x_ = 3 [pixels] is the size of the phase processing windows used. Inevitably, the application of phase-processing windows in both cross-correlations resulted in the axial and lateral resolutions of the strain maps being reduced to 39 and 144 μm, respectively, which is lower than the original resolution of the structural images.

## Appendix C

To exhaustively describe the hyperleastic material model we reported in the method section, we have to start from eq. (1) of this manuscript. *Ī*_1_ and *Ī*_4_ are invariants of the isochoric Cauchy-Green strain tensor. 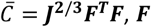 is the deformation gradient tensor, and *J* = det (***F***). The Macaulay bracket operator, denoted as < • >, is employed to account for the fact that collagen fibers only contribute to the overall mechanical response when they are in a state of tension:

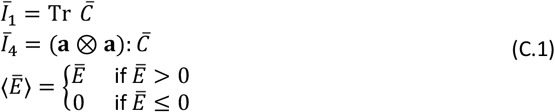

The general fiber direction in spherical coordinates, ***a***, is described as:

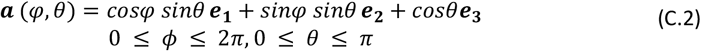

The angular density of the fiber distribution ρ was decomposed as a product of the in-plane and out-of-plane distributions *ρ*_*in*_ and *ρ*_*out*_:

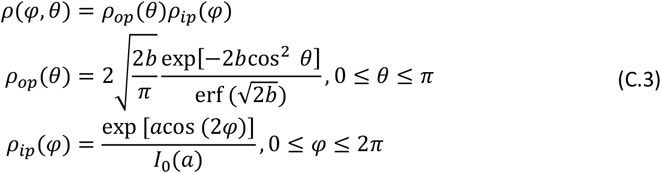

Where, *I*_0_ is the modified Bessel function of the first kind of order 0. The fiber distribution parameters *a* and *b* were defined based on x-ray scattering and second harmonic generation data. While *b*=2.5 was kept constant along the corneal curvature, the parameter *a* went from *a*=0 (isotropic in-plane dispersion) in the anterior part to *a*=5 (aligned fibers direction) in the posterior part of the cornea.

## Appendix D

This study employed two different meshing approaches to create a FEM of the cornea (Figure D1). For identifying *ex vivo* mechanical parameters, an average cornea model was used to reduce computational costs, assuming corneal geometry has minimal effect on NI experiments. When applied to the *in vivo* clinical setting, however, a patient-specific approach was adopted to generate meshes tailored to individual corneal geometries, ensuring accurate curvature analysis in post-CXL evaluation.

**Figure D1:**
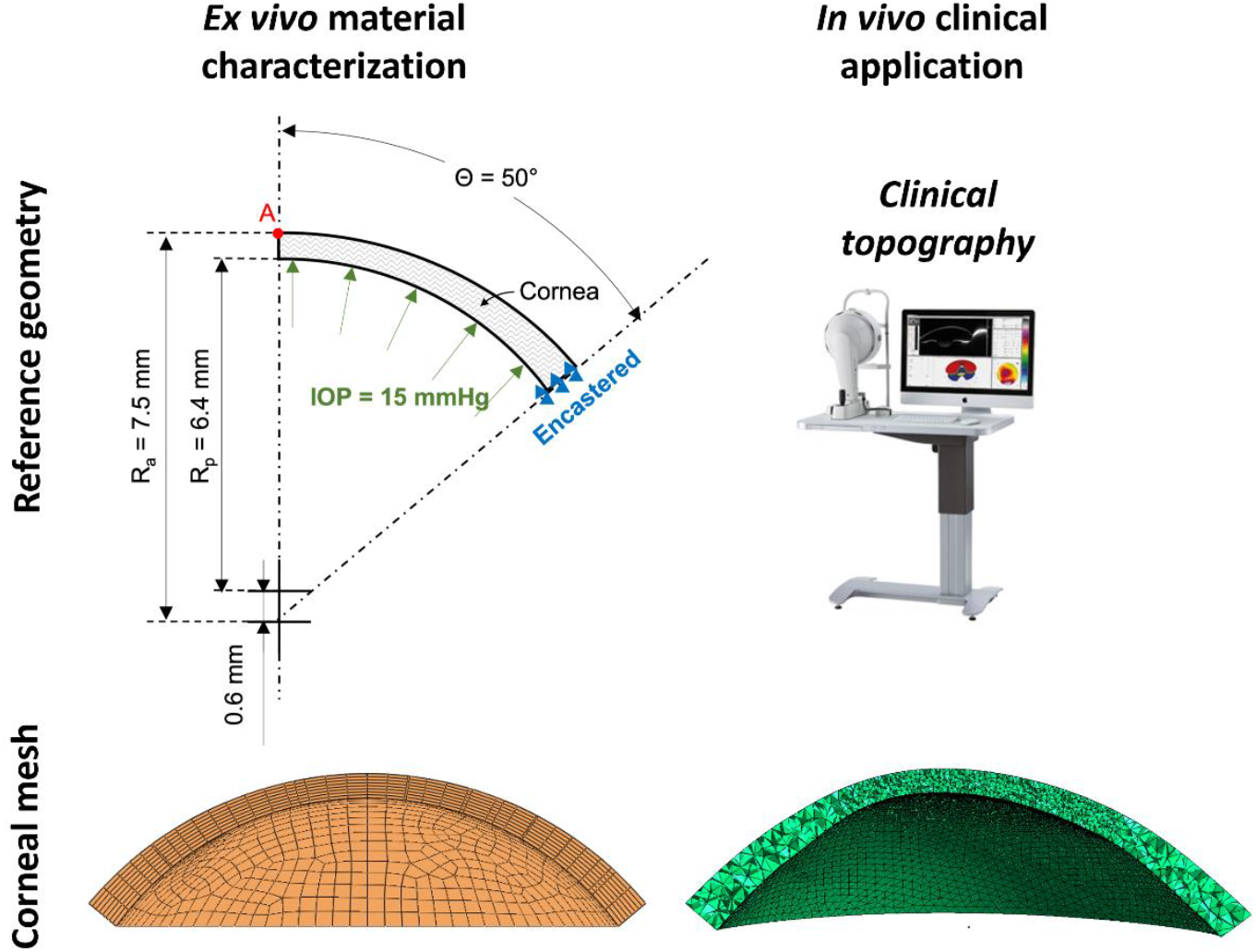
different corneal reference geometries and corresponding meshes adopted in the different phases of the present study.

## Appendix E

## Methods

### Uniaxial tensile test

Strips measuring 6×2 mm were cut from both the CXL-treated and non-CXL portions of the cornea using the Femto LDV Z8 Neo femtosecond laser (Ziemer Ophthalmic Systems AG, Switzerland). These strips were then preserved overnight in a MEM 15% dextran solution. Each strip was subjected to uniaxial testing using the UStretch device (CellScale, Waterloo, Canada). The tests were conducted at room temperature with the strips maintained in the culture medium bath. Each strip was pre-stretched with a force of 10 mN and subjected to 6 loading cycles at a strain rate of 0.16%/s, each reaching a strain of 10%. The force-displacement data from the fifth cycle were analyzed, and the tangential elastic modulus at 10% strain (*E*_10_) was considered for comparing the stiffness of the UV irradiated regions vs non-irradiated ones.

### Results

#### Uniaxial tensile test

Figure E1 shows that the corneal tissue exhibited a stiffer, but statistically non-significant behavior under UTT after CXL, which was reflected by an increased tangent modulus (124±77 kPa vs 164±70 kPa; p=0.125). Out of n=5 corneas considered, one was discarded since one of the strips got damaged during the mounting phase.

**Figure E1:**
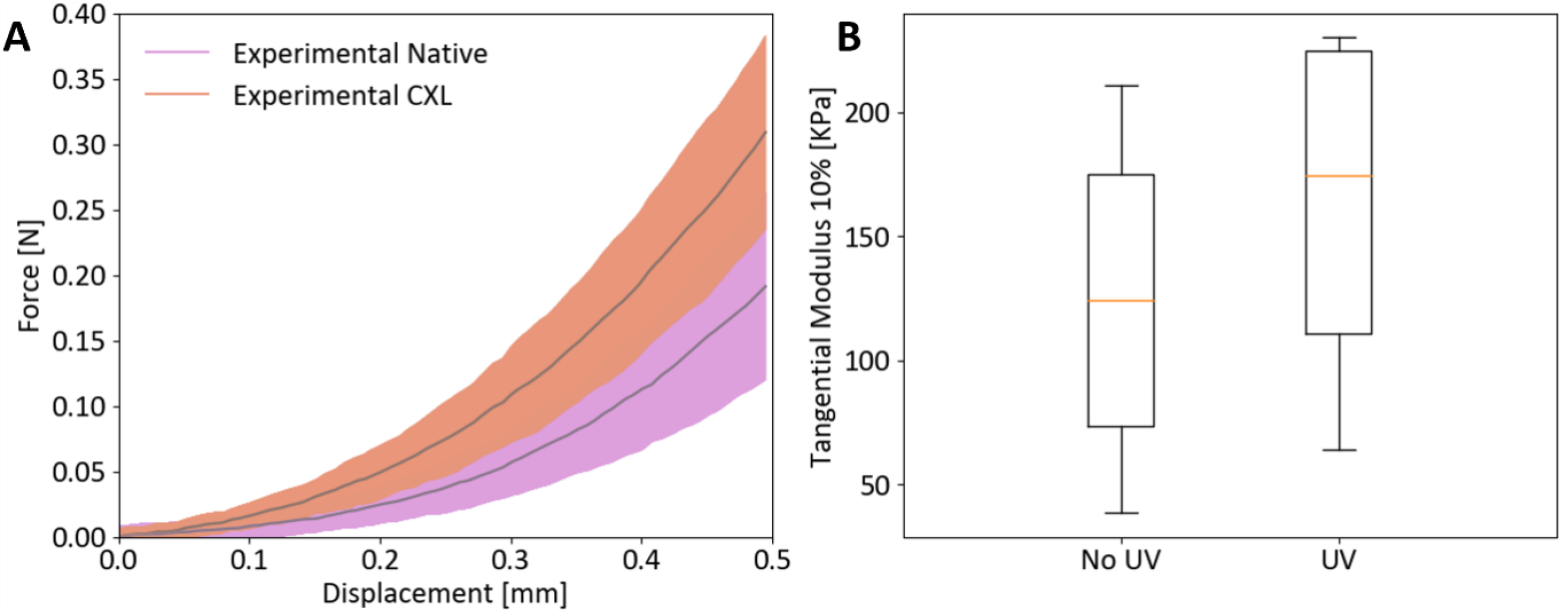
no-UV vs UV region UTT tensile test results (n=4). A) experimental force displacement curves. B) tangential modulus at 10% strain distributions.

#### Nanoindentation

As shown in Figure E2, the non-irradiated region reported a slightly higher *E*_*HZ*_ when compared to the irradiated one after CXL, with values ranging from 145.7±7.5 kPa to 140.9±4.7 kPa (p<0.01), respectively. The decrease in *E*_*HZ*_ here reported (4.75±9.6 kPa) is in the range of the standard deviations of the NI measurements performed in this work. No statistically significant variations in *C*_*IT*_ were reported between these two areas of the cornea (16.23±1.4 % vs 16.51±1.3 %, p=0.09, for non-UV and UV regions, respectively).

**Figure E2:**
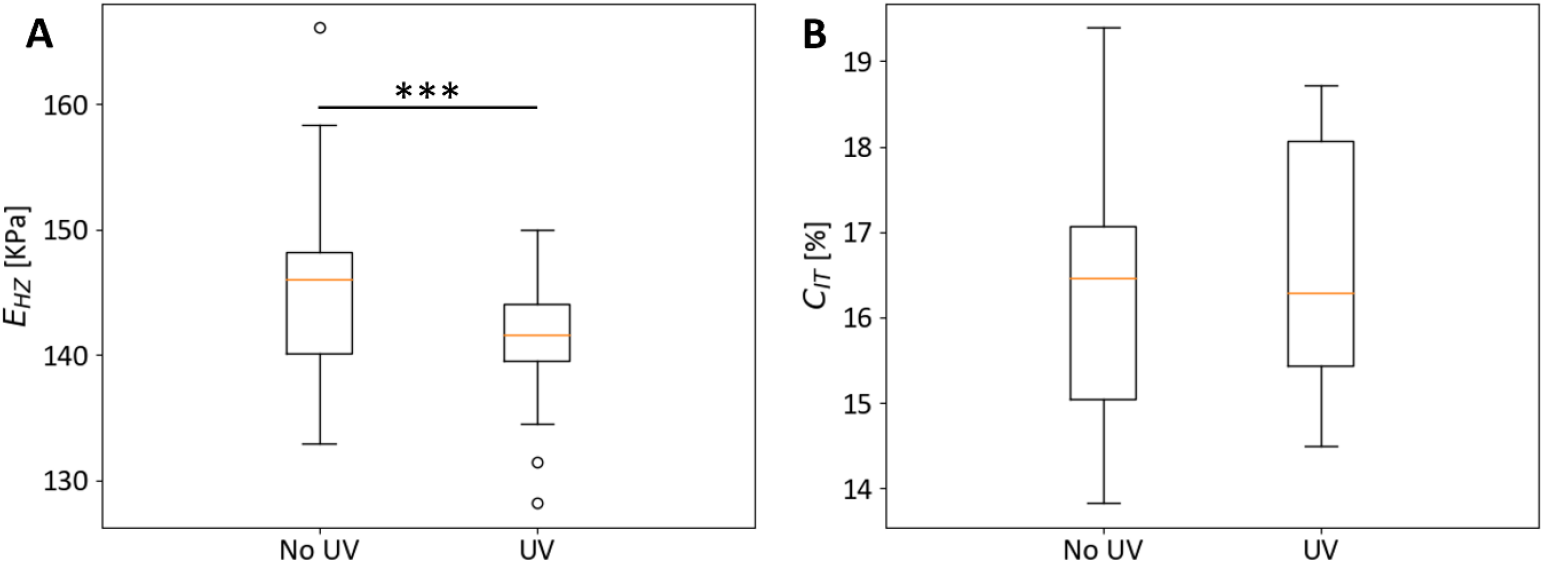
no-UV vs UV region NI test results (n=25). A) Elastic Hertz modulus *E*_*HZ*_. B) indentation creep *C*_*IT*_. ***.001≤p≤.01.

#### Ex vivo optical coherence elastography

No statistically significant differences were observed between the non-irradiated and the irradiated regions in terms of *ε*_*zz*_ measured via *ex vivo* OCE (0.5±3.1 ‰ vs 1.1±2.6 ‰, p=0.125, for the no-UV and UV regions, respectively), as reported in Figure E3. From the enface view in Figure E4 the patterned CXL effect of the FEM can be appreciated.

**Figure E3:**
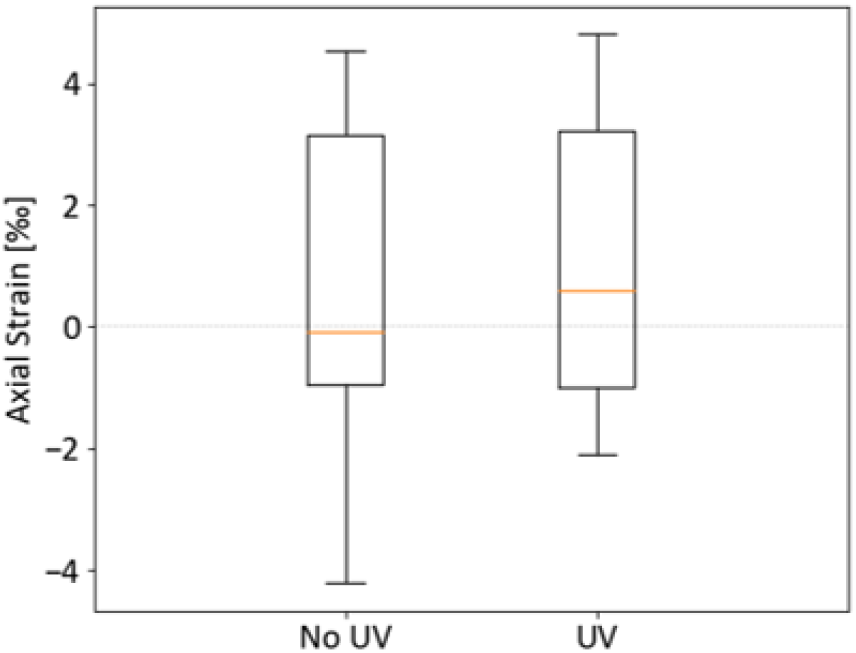
no-UV vs UV region *ex vivo* OCE inflation test results (n=5).

**Figure E4:**
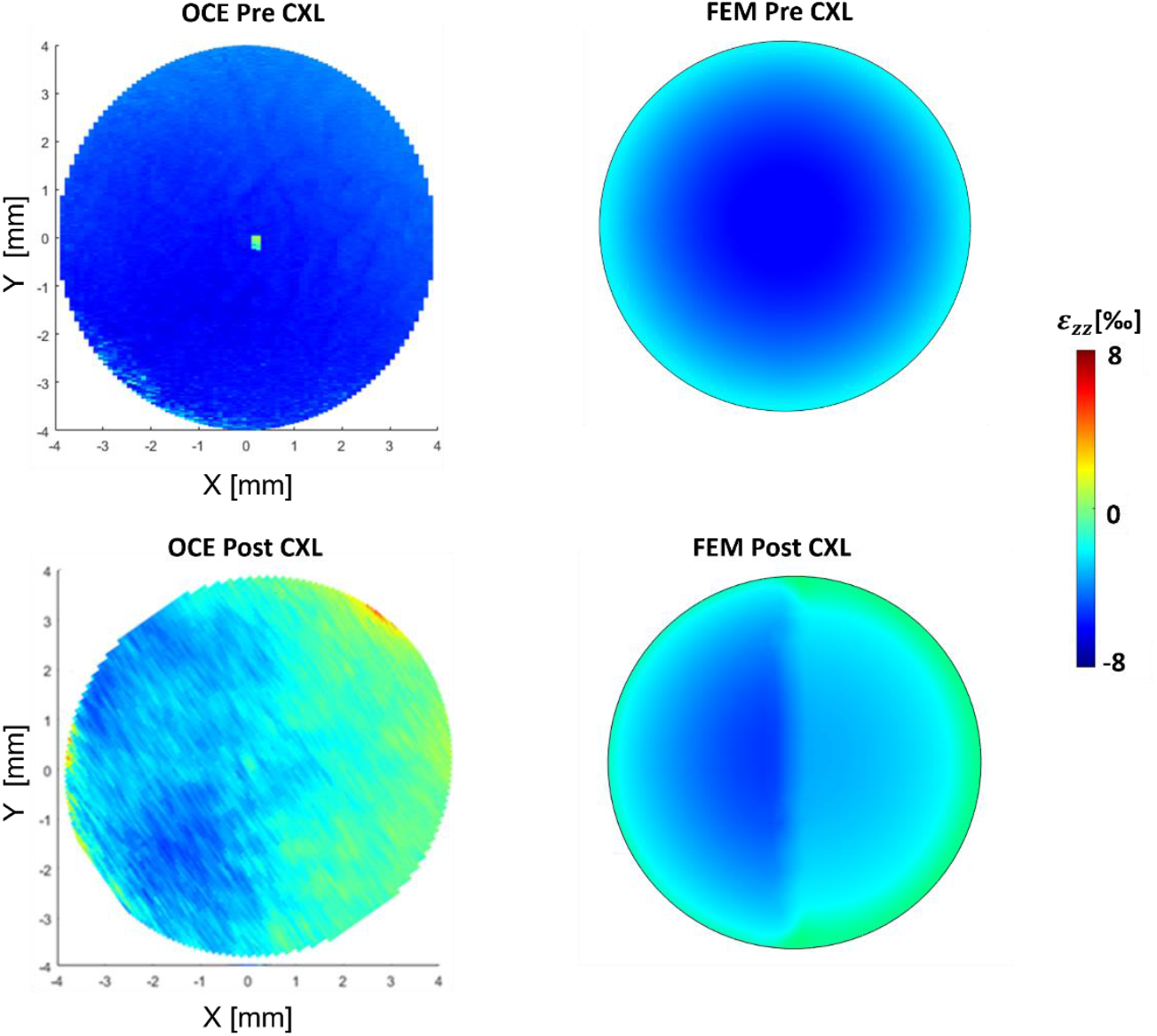
enface views for both OCE imaging (left column) and FE results (right column). The first row shows the pre-CXL condition, while the second row shows the post-CXL condition. Color scale represents axial strain *ε*_*zz*_ [‰] values.

## Notes

### Competing Interest Statement

The authors have declared no competing interest.

